# On the optimal design of metabolic RNA labeling experiments

**DOI:** 10.1101/428862

**Authors:** Alexey Uvarovskii, Isabel S. Naarmann-de Vries, Christoph Dieterich

## Abstract

Massively parallel RNA sequencing (RNA-seq) in combination with metabolic labeling has become the *de facto* standard approach to study alterations in RNA transcription, processing or decay. Regardless of advances in the experimental protocols and techniques, every experimentalist needs to specify the key aspects of experimental design: For example, which protocol should be used (biochemical separation vs. nucleotide conversion) and what is the optimal labeling time? In this work, we provide approximate answers to these questions using asymptotic theory of optimal design. Specifically, we derive the optimal labeling time for any given degradation rate and show that sub-optimal time points yield better rate estimates if they precede the optimal time point. Subsequently, we show that an increase in sample numbers should be preferred over an increase in sequencing depth. Lastly, we provide some guidance on use cases when laborious biochemical separation outcompetes recent nucleotide conversion based methods (such as SLAMseq).

## Introduction

Changes in gene expression are frequently observed in pathological conditions. In the simplest model [Schwanhäusser et al., 2011], steady state RNA levels are governed by synthesis (transcription) and degradation rates (RNA stability). A paradigm is the generation of the hypoxic response in pathological conditions such as heart insufficiency [Ziaeian and Fonarow, 2016] and fast growing tumors [Wilson and Hay, 2011]. Hypoxia (< 2% O_2_) results in a global decrease of total transcription [Johnson et al., 2008]. However, transcription of specific target genes is induced under hypoxic conditions by hypoxia inducible factor 1 (HIF1) [Semenza, 2003], which is composed of a stable *β*-subunit and an oxygen labile *α*-subunit [Huang et al., 1998]. Furthermore, different RNA binding proteins such as HuR and TTP as well as miRNAs regulate the stability of their cognate target mRNAs dependent on oxygen availability [Gorospe et al., 2011] and contribute to changes in gene expression profiles.

Metabolic labeling experiments are a versatile tool to discern dynamic aspects in physiological and pathological processes. These experiments drive our understanding of key processes in molecular systems, such as synthesis and decay of metabolites, DNA, RNA and proteins. Pulse-chase experiments help to determine the kinetic parameters of synthesis and decay in various contexts. In the pulse phase of an experiment, the label is introduced to newly synthesized compounds and unlabeled or pre-existing molecules are only subjected to degradation or some other form of processing. In contrast, during the chase phase, the label in the system is gradually replaced by unlabeled compounds. A typical metabolic labeling experiment may include a pulse, a chase or both phases.

The first transcriptome-wide studies by Cleary et al. [2005] and Dölken et al. [2008] used 4-thiouridine (4sU) labeling in cell culture experiments to infer kinetic parameters. This approach has become quite popular in RNA biology, which is shown by a vastly increasing number of studies (see Wachutka and Gagneur [2017] for review).

Massively parallel RNA sequencing (RNA-seq) in combination with metabolic labeling has become the *de facto* standard approach to study alterations in RNA transcription, processing or decay at the transcriptome-wide level. At the time of writing, the most widely used approach involves metabolic labeling with thiol-labeled nucleoside analogs such as 4sU (4sU-tagging) [Baptista and Dölken, 2018]. Briefly, total cellular RNA is isolated and thiol groups are biotinylated. Subsequently, total cellular RNA can be efficiently separated into newly transcribed (labeled) and pre-existing (unlabeled) RNA.

A very recent achievement is the arrival of new methods involving chemical conversion of 4sU residues into cytosine analogs, which is observed as point mutations in RNA-seq data (T-to-C transitions), (see Herzog et al. [2017], Schofield et al. [2018] and Riml et al. [2017]). The absence of any biochemical separation method makes metabolic labeling more accessible due to lower input amounts and less laborious protocols.

Regardless of advances in the experimental protocols and techniques, a few important questions remain to be answered by any experimentalist, namely the specifics of experimental design: What should be measured (i.e. sequenced) and when? For example, which approach should I take (e.g. biochemical separation vs. nucleotide conversion), when should I collect my samples (e.g. time points in a pulse experiment) and how could this affect my estimates on kinetic parameters.

Within this manuscript, we use kinetic and statistical models to infer degradation rates from a pulse experiment (see Figure 1 and Equations 1-2), and derive several aspects on the optimal design of metabolic RNA labeling experiments.

**Figure 1:**
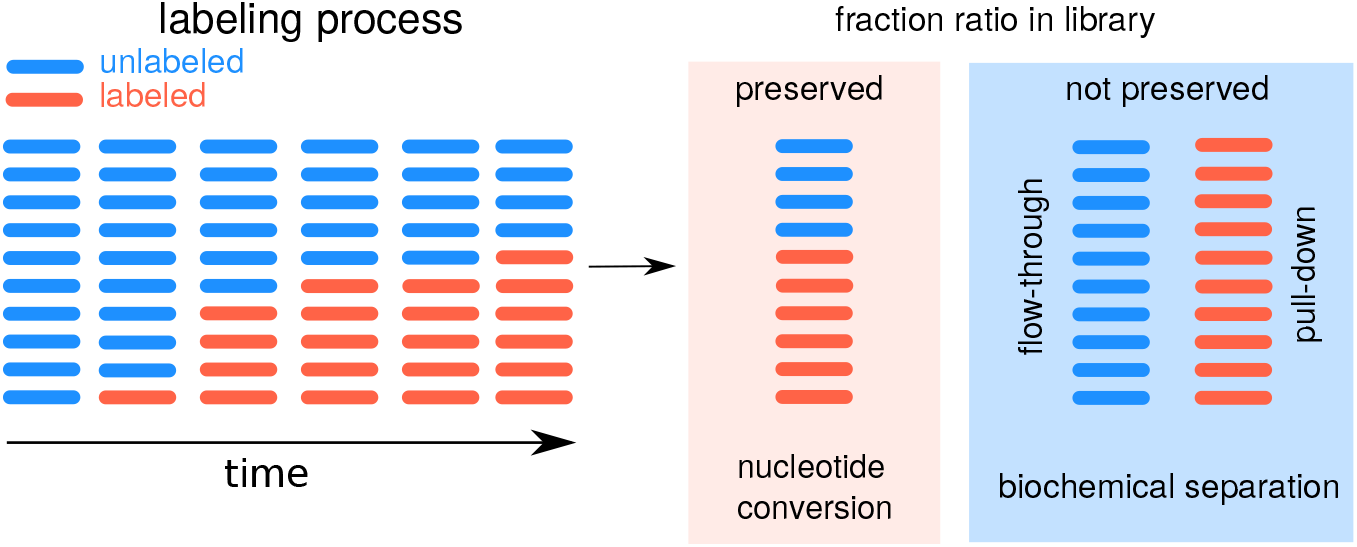
Pulse labeling experiment types to measure degradation rates. The conventional approach as in Duffy et al. [2015] utilizes biochemical separation, which does not preserve the fraction ratio (labeled vs. unlabeled) in the read counts. Alternative novel approaches (e.g. Herzog et al. [2017]) induce reverse transcription signature events (nucleotide conversions, typically T-to-C). Individual reads can be classified by the presence or absence of this characteristic nucleotide conversions. For this particular case, the fraction ratio is well reflected by the read counts.

## Results

### Model of experiment

We describe RNA-seq read counts with the negative binomial distribution, which is widely used in this setting and accounts for overdispersion [Anders and Huber, 2012]. For a given gene, read count follows *X ~ NB*(*m*(*μ, d, t*)*, k*), where *m* is the mean read count, which depends on the time of labeling *t*, the degradation rate *d* and the expression level in the steady-state *μ*, and *k* is the overdispersion parameter of the negative binomial distribution *NB*. In this case, variance var(*X*) = *m*(*m* + *k*)*/k*, where low *k* values correspond to high overdispersion in the data.

We describe RNA amount in the labeling experiments using simple first order kinetics:

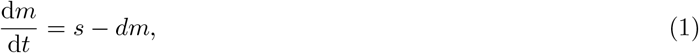

where *s* is the synthesis rate and *d* is the degradation rate. In a steady-state, the expression level of a gene is *μ* = *s/d*. The expression level *μ* can be derived from the total fraction, which ensures identifiability of at least this parameter. For that reason, we use *μ* and *d* to parametrize the model. In this manuscript, we only discuss the case of pulse labeling experiments throughout. However, our considerations extend to chase labeling experiments, where equations are the same, except that the labeled fraction behaves as the unlabeled one in the pulse experiment and *vice versa*. For simplicity, we assume that fraction cross-contamination is negligible, in which case, RNA amounts for a given gene are proportional to the means *m_L_*, *m_U_* and *m_T_* derived from the kinetics for labeled, unlabeled and total fractions scaled by sample-specific factors *x*_i_:

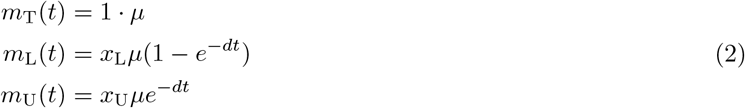

Here we treat mean read count in the total sample as a reference (coefficient is 1), to make the system identifiable. In the case of labeled and unlabeled fractions, expected read numbers must be scaled by additional coefficients, *x*_U_ and *x*_L_, because RNA material can be normalized by different degrees during library preparation from chemically separated fractions.

Certain experimental approaches preserve the ratio of labeled to unlabeled fractions (see Figure 1; e.g. SLAMseq), *x*_U_ = *x*_L_, because they do not involve any biochemical purification step. If the sequencing depth is approximately the same for all samples, we may assume for simplicity *x*_U_ = *x*_L_ = 1, and in this case, *m*_T_ = *m*_L_ + *m*_U_ = *μ*.

In the conventional approach, where labeled and unlabeled molecules are separated, *x*_U_ = *x*_L_, and the fraction ratio must be inferred from the data itself or by using an external normalization by spiking in labeled and unlabeled known molecules. In the presence of cross-contamination, the estimations for the rates are biased depending on the relation of the labeling time and the degradation rate: if *dt* ≪ 1 (slow rate), the bias is towards faster rate values, and, if *dt* ≫ 1 (fast rate), it is towards slower rate values, see Extended methods for more details.

### Best time to measure

To infer the values of model parameters ***θ*** (where elements of the vector *θ_i_* correspond to *μ*, *d* etc.), we maximize the likelihood function *ℒ* (***θ****, X*) given read counts *X*.

In this paper, we derive our results on the basis of the asymptotic theory, when the number of experiment repetitions *n* →∞, in which case the system can be treated analytically. However, it provides only an approximation, which depends on how close the log-likelihood is to its quadratic approximation. In this case, for *n* →∞ and under a number of regularity conditions, the maximum likelihood estimator (MLE) ***θ̂*** is consistent and its covariance matrix can be approximated by the expected Fisher information matrix (FIM)

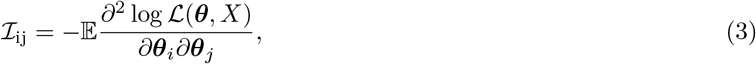

then distribution of ***θ̂*** is asymptotically normal:

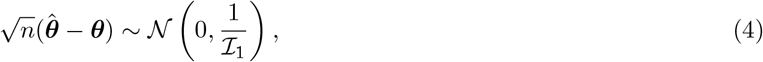

where *ℐ*_1_ corresponds to the FIM for a single experiment repetition [Chernoff, 1953, Pawitan, 2001]. We assume that the overdisperion parameter *k* is shared between all genes and neglect the uncertainty in *d* propagating from *k*, i.e. only two parameters, *d* and *μ*, are used to construct the FIM:

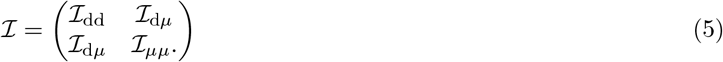

The FIM is additive, i.e. if *ℐ*_U_ and *ℐ*_L_ correspond to the labeled and unlabeled fractions, the total FIM for the experiment is *ℐ* = *ℐ*_U_ + *ℐ*_L_

The diagonal terms of the inverse FIM estimate the variance of *θ̂_i_*

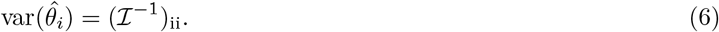

Since the FIM *ℐ* depends on the experiment parameters, such as labeling time *t* and sequencing depth, it is our main interest is to reduce variance of the MLE by selecting the optimal conditions accordingly.

Noting that *ℐ*= *nℐ*_1_, where *ℐ*_1_ is the FIM for a single measurement, we will omit *n* and will optimize the FIM for *n* = 1 instead.

In the case of multiple parameters, it may be not possible to achieve minimal variance for all parameters at the same time. Different criteria can be constructed as a combination of the elements of the inverse FIM [Chernoff, 1953, Van den Bos, 2007]. We are interested to optimize estimation of *d* only and do not consider variance of the expression level estimator *μ̂* in the design criteria.

Let us consider first a simpler experimental setup, which preserves the fraction ratio (e.g. SLAMseq). The derivations for a general case are left to the Extended Methods section, and here we first discuss the case of the Poisson model, which corresponds to the case of no overdispersion (*k* →∞). Let *X*_L_ and *X*_U_ be the read counts corresponding to the labeled and unlabeled molecules for a given gene in a SLAM-seq sample, and let *t* be the time of labeling. In this case, the inverse FIM is diagonal:

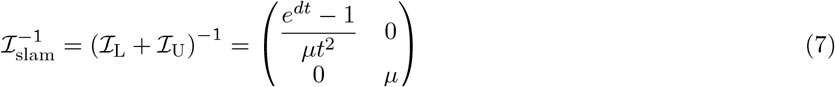

The parameters *d* and *μ* are information orthogonal, because *ℐ*_d*μ*_= 0 and inference about *d* can be done as *μ* were known exactly.

Indeed, for *X*_L_ ~ Pois(*m*_L_), *X*_U_ ~ Pois(*m*_U_), the conditional distributions *P* (*X*_L_ *X*_U_ + *X*_L_) and *P* (*X*_U_ *X*_U_ + *X*_L_) are binomial with the rates *m*_U_*/*(*m*_U_ + *m*_L_) = *e^−dt^* and *m*_L_*/*(*m*_U_ + *m*_L_) = 1 − *e^−dt^* and do not depend on *μ*. This model was recently discussed in a Bayesian framework for SLAMseq experiments by [Jürges et al., 2018].

For a diagonal *ℐ*, the inverse term 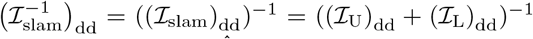. The maximum of the term (*ℐ*_slam_)_dd_ corresponds to minimal asymptotic variance of *d*̂ due to Equation 4. By optimizing (*ℐ*_slam_)_dd_ in respect to *t*, we get

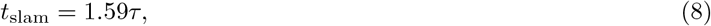

where *τ* = 1*/d* is the characteristic time of degradation. That means, that if one optimizes the SLAMseq experiment and is concerned with characteristic time of degradation of *τ*, the measurement at time point 1.59*τ* corresponds to the asymptotically optimal design. For example, if one is interested in an RNA species with half-life time of *λ* = 1 hr (i.e. the characteristic time *τ* = *λ/* log(2) ≈ 1.44 hr), a measurement at around 1.59 × 1.44 ≈ 2.3 hr corresponds to the asymptotically optimal design.

In Figure 2A, we depicted dependency of (*ℐ*_slam_)_dd_ and corresponding values of (*ℐ*_U_)_dd_ and (v_L_)_dd_ as functions of normalized time *t/τ* for the degradation rate *d* = 1.

**Figure 2:**
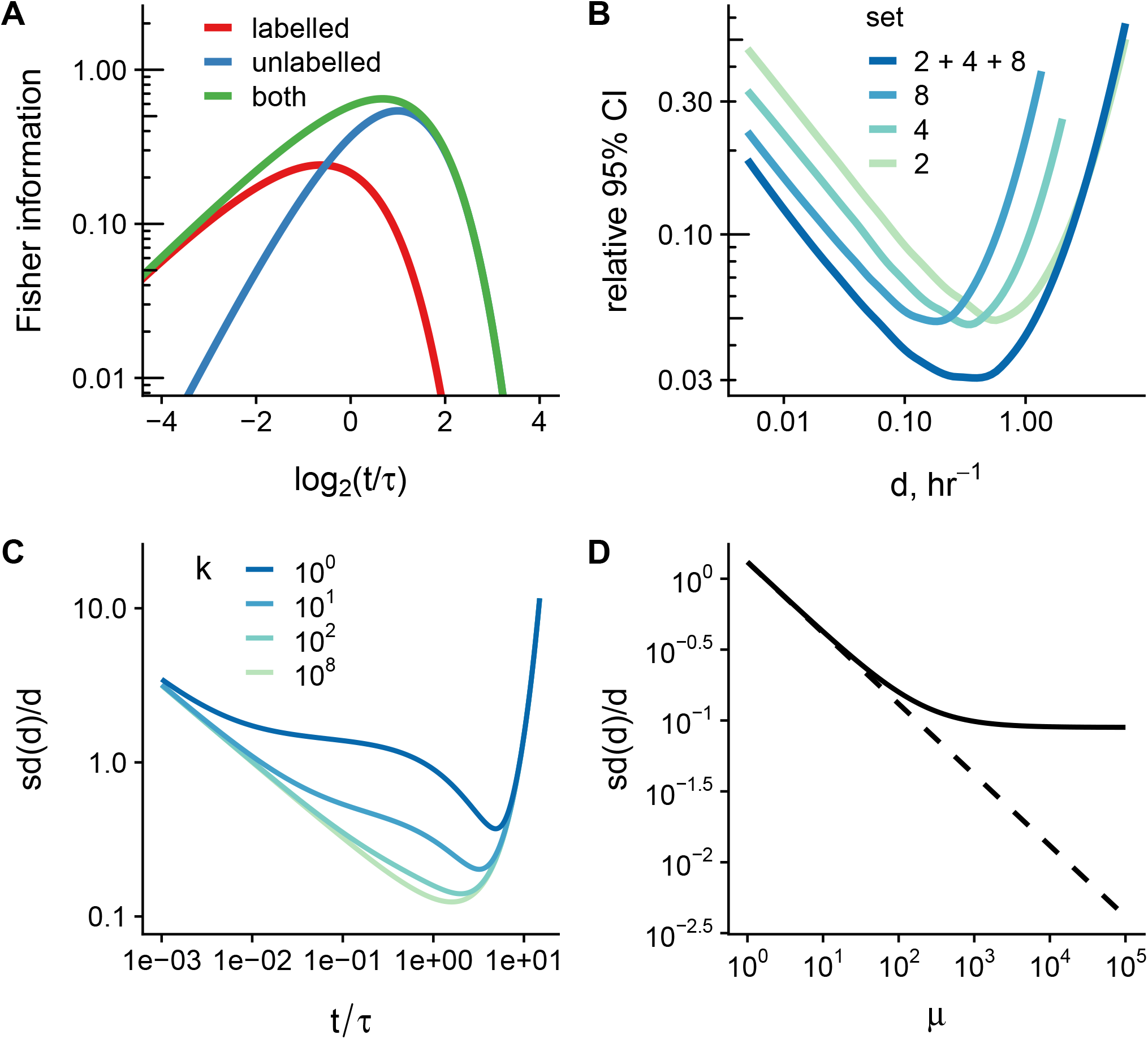
Key characteristics of metabolic RNA labeling experiments. **A:** The diagonal term of the Fisher information matrix (FIM) *ℐ*_dd_, which corresponds to the degradation rate, as a function of the ratio of labeling time to the characteristic time of degradation *τ* = 1*/d* for the case of SLAMseq experiment. Read counts follow the Poisson distribution, the expression level is *μ* = 1. **B:** 95% confidence interval (CI) relative width of the degradation rates for different sets of time points included in the simulation of the SLAMseq experiment. We simulated counts for a range of rates *d* and assumed for simplicity that normalization factors are perfectly known but not the rates and expression levels. Smoothed data from 10 simulation runs is shown. **C:** Relative standard deviation (*sd*(*d̂*)*/d*) of the MLE estimator for *d* as a function of measurement time at different values of the overdispersion parameter *k*. With increasing overdispersion, the profile of the dependency flattens. However, near the optimal time point, variance of the estimation is more sensitive to time of labeling, which complicates the optimal design choice for different *d* ranges. Expression level is fixed to *μ* = 100 reads in this example, the degradation rate is assumed to be *d* = 1. The FIM *ℐ* = *nℐ*_1_ is calculated for *n* = 1. **D:** Relative standard deviation (*sd*(*d̂*)*/d*) for a model with overdispersion (*k* = 100, solid line) or with no overdispersion (*k → ∞*, dashed line). The degradation rate is *d* = 1, the labeling time is *t* = 1. The FIM *ℐ* = *nℐ*_1_ is calculated for *n* = 1.

Interestingly, (*ℐ*_U_)_dd_ and (*ℐ*_L_)_dd_ achieve maximum at *t_U_* = 2*τ* and *t_L_* = 0.64*τ*, and the main contribution to the sum (*ℐ*_slam_)_dd_ = (*ℐ*_U_)_dd_ + (*ℐ*_L_)_dd_ comes from the term corresponding to labeled counts at shorter labeling times, and from the term for unlabeled counts at times longer than *τ*, see Figure 2A.

### Too early is better than too late

Usually one is interested to measure a rate with a certain relative precision. To reflect this, we normalize the variance of the estimator by *d*^2^:

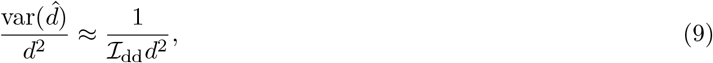

Technically, such modification corresponds to the transformation *d* = *e^η^*, because 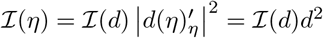, as if the degradation rate were considered at the logarithmic scale, and any absolute changes in *η* would correspond to relative changes in *d*. However, we avoid introducing additional parameters and stick to the usage of *d* only.

For brevity we omit the index, since we consider only the *dd* diagonal term of *ℐ* in this subsection. By introducing a non-dimensional time variable *α* = *t/τ*,

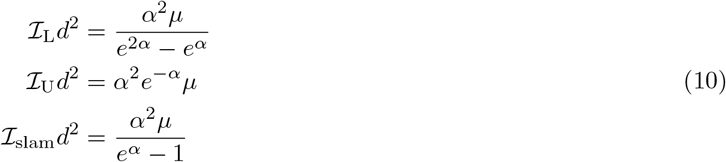

For labeling times much shorter than characteristic degradation time of a given gene, *α* ≪ 1, and the normalized FIM terms behave as a power function:

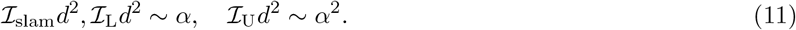

However, for labeling times much longer than the characteristic time of degradation *τ*, *α*≫1, the normalized FIM terms vanish exponentially:

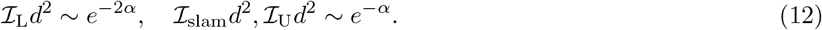

In a typical high-throughput experiments, kinetic parameters are monitored for a large set of genes (in the order of thousands), which may have different degradation rates. In this case, every time point in the experiment will be only optimal for a subset of these genes. To illustrate this effect, we simulated read counts for a SLAMseq experiment scheme (with no over dispersion) and fitted the model using various sets of samples. In our *in silico* experiment, we always included the total fraction (*t* = 0 hr), and either one additional time point (labeled and unlabeled fractions) or all time points (2, 4, and 8 hr). The normalization coefficients were set to 1 to mimic the SLAMseq scheme, as discussed earlier, Equation 2.

We fitted the model using the pulseR package and computed the 95% confidence intervals (CI) for *d* using the profile likelihood approach [Uvarovskii and Dieterich, 2017]. Since we assume no overdispersion (Poisson distribution), for high read counts (*μ* = 10000) the quadratic approximation of the log-likelihood function applies, and the confidence intervals for the rate estimations may be approximated by the Wald intervals, i.e. 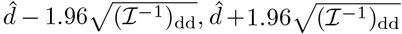, and hence, they reflect the behavior of the FIM term for *d*. As expected, the relative CI width is minimal only for a certain subset of the rates, depending on the set of measurements included, see (Figure 2B).

If the degradation rate is very fast in comparison to the experiment time scale, the CI width for these fast genes is defined by the earliest time point in the experiment (see Figure 2B).

Since every labeling time is optimal only for a single degradation rate, it might be beneficial to focus the design on genes with faster rates *d*, if sample size is limited and no other criteria of optimality are given. The justification follows from the faster decay of the FIM term for *α* ≫ 1 (i.e. genes with faster kinetics), Equations 11, 12.

### Increasing sample numbers is preferred over higher sequencing depth

Distribution of read count from RNA-seq experiments exhibits overdispersion, and negative binomial distribution (NB) can be used to account for that [Anders and Huber, 2012]. In this section, we explore how presence of overdispersion would affect inference about *d*. The overdispersion parameter *k* of the NB distribution describes the level of overdispersion in the data, in which case the variance is defined as var(*X*) = *m* + *m*^2^*/k* for counts *X* ~ *NB*(*m, k*) with mean *m*. Smaller values of *k* correspond to higher overdispersion level, and, for *k*→∞, the NB distribution converges to the Poisson distribution, for which var(*X*) = *m*. For simplicity, we assume that distributions of read counts in all samples share the same value of *k*. In addition, we do not consider uncertainty in the overdispersion parameter *k* when we make inference about *d* for individual genes, in a way as it is implemented in some packages for differential expression analysis, for example, in DESeq, [Anders and Huber, 2012]. A more advanced quasi-likelihood approach, which accounts for uncertainty in the overdispersion parameter, is discussed in Lund et al. [2012].

In the case of NB distribution, the FIM is not diagonal for the SLAM-seq experiment, see the Extended Methods section. Hence we need to work with the inverse FIM, and the diagonal term for the SLAMseq design is

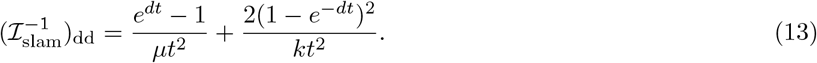

The presence of overdispersion shifts the optimal time to higher values. But the most important change is that the profile of 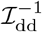 is more sensitive to the labeling time *t* near the optimal point. For higher overdispersion values, the variance of the rate estimator *d̂* increases faster in the vicinity of the optimum (see Figure 2C). It imposes stricter conditions on the experiment design. The second term in the Equation 13 vanishes for times *t* ≫ 1, and the equation coincides with the case of no overdispersion. The contribution of the second term is higher for smaller values of *k* (higher overdispersion) and for shorter labeling times *t*, with maximal value at *t →* 0:

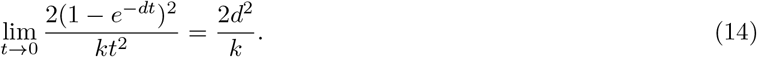

Another limitation, which arises in the over-dispersed model is that increase of the sequencing depth has a limiting effect on the variance. Indeed, only the first term in Equation 13 can be eliminated by increase of sequencing depth:

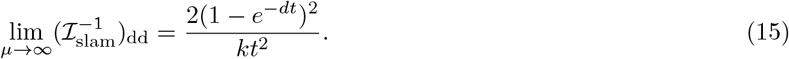

In contrast, repeating the experiment *n* times affects both terms in 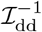, since for *n* repetitions,

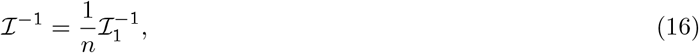

where 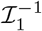 is the inverse FIM for one repetition. That means, that it can be more beneficial to spread the sequencing capacity between several biological replicates. In the limiting case *n* →∞, if the total depth of *n* repetitions is fixed to the initial value such, that *nμ*_1_ = *μ*, where *μ*_1_ corresponds to the reduced depth in a single repetition,

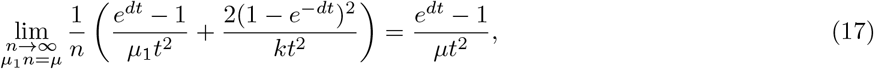

which coincides with the FIM term for the Poissonian (non-overdispersed) model, compare to Equation 7.

In the Poissonian case, when *k* →∞ and the second term is absent (see Equation 7), adding twice more samples or increasing sequencing depth by two fold results to the same FIM and, consequently, the same approximation of the variance var(*d̂*). In the logarithmic scale, the Poissonian case is represented by a line, Figure 2D (dashed line), which can be compared to the case of overdispersion (solid line), when an increase in the depth is inefficient starting from a certain value. A similar phenomenon was discussed by [Robles et al., 2012] in the context of differential gene expression analysis by RNA-seq.

### Biochemical separation still matters

If one is interested in estimating rates of extreme values by using very short (e.g. TT-seq, Schwalb et al. [2016]) or long labeling times, it may be less efficient to use the protocols, which preserve the ratio of labeled and unlabeled molecules (e.g. SLAMseq). Let us consider a study of fast gene kinetics, in which case, very short labeling times are used. In this case, *dt* ≪ 1 for majority of the genes, and the labeled fraction constitutes only a minor proportion of the input SLAMseq sample, because *m*_L_ = *μ*(1 − *e^−dt^*) ≈ *μdt* ≪ 1. The input SLAMseq sample mainly contains unlabeled molecules from the non-target slower genes, which results into spending sequencing resources on non-informative material. The same idea holds for very long times, when *dt* ≫ 1 and most of unlabeled molecules are subjected to degradation, *m*_U_ = *μe^−dt^* ≪ 1.

In contrast, conventional experimental setups with a separation step can be used to focus sequencing capacity on relevant molecules. However, the conventional approach suffers from the need to normalize sequencing results from different fractions as it does not preserve the ground truth ratio of labelled and unlabelled molecules. In typical RNA-seq experiments, normalization coefficients assumed to be shared between all the genes in a given sample [Anders and Huber, 2012], but nevertheless, it introduces additional uncertainty into rate estimations and further analysis is required to estimate level of this contribution. In the following derivations, we neglect the uncertainty in estimating the fraction normalization coefficients *x_i_* from the Equation 2.

To illustrate the benefit of the conventional approach, let us consider a set of fast genes, such that there exists labeling time, when majority of genes *i* ∉ *ℱ* do not contribute to the labeled fractions, i.e 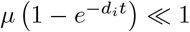 for *i* ∉ *ℱ*, but *μ* 1 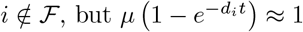 for *i ∉ ℱ*. If the sequencing depth of the labeled fraction is approximately the same as for the total sample, then the normalization factor is

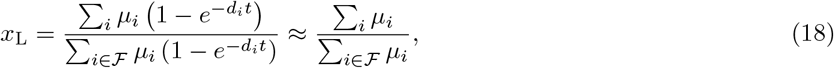

which can be high at short times. Such “zooming” effect can be considered as corresponding increase of the sequencing depth in the SLAMseq experiment by the factor of *x*_L_ for the labeled fraction. The same idea can be applied to the unlabeled fraction and long labeling times, when the sequencing depth is shared out between the most stable set of genes. Since the normalization factor depends on the rate distribution and expression level in a given system, it is not possible to derive optimal design criteria analytically without imposing additional assumptions.

Since

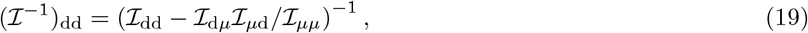

and using the fact that *ℐ*_d*μ*_= *ℐ_μ_*_d_ and *ℐ_μμ_ >* 0, the diagonal term of the inverse matrix is bounded

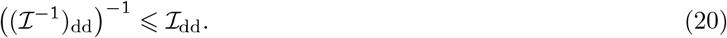

The diagonal element of the FIM *ℐ*_dd_ represents the upper bound of the information gain in a given design, corresponding to assumption of no uncertainty in parameters other than *d*. As in the case of the SLAMseq, inference can be improved to a limited extent by increase of sequencing depth, if overdispersion is present in the data, compare to Equation 15:

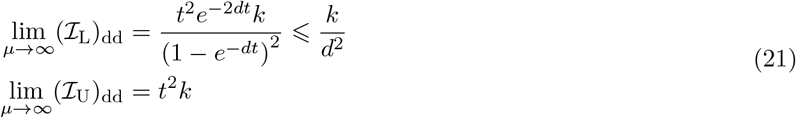

It is interesting to note, that for the case of the unlabeled fraction, the bound can be improved by use of longer labeling times (provided very high sequencing depth), which is not the case for the labeled fraction (with the upper bound *ℐ*_L_ *→ k/d*^2^ at *t →* 0).

In summary, biochemical separation should be considered for estimation of degradation rates of RNA species with extreme values. Another design choice is to reduce the number of sequencing reactions by using external spike-ins. For slowly turned over RNA species, one may sequence total and unlabeled fractions, and, for fast turned over RNA species, the total and the labeled fractions. The use of external spike-ins ensures identifiability of the normalizing coefficient from only two fractions.

### Example from a pulse labeling experiment

MCF-7 cells were pulse labeled with 200 *μ*M 4sU for 2, 4 or 8 hrs. 4sU-labeled and unlabeled RNA were separated by streptavidin purification after MTSEA biotin-XX catalyzed biotinylation of 4sU-labeled RNA [Duffy et al., 2015]. The efficiency of purification was monitored in a dot blot assay that detects biotinylated RNA with streptavidin-HRP (Figure 3A). This analysis revealed a gradual increase in biotinylation with increasing labeling time. Importantly, biotinylated transcripts were efficiently depleted from the flow trough, as no biotinylation signal could be detected in these samples. Biotin-enriched RNAs are eluted by de-biotinylation with DTT. Therefore, we estimated purification efficiency by the amount of purified RNA determined by A_260nm_ absorption measurement. The amount of purified RNA increased gradually with increasing labeling time (Figure 3B) comparable to the biotinylation signal increase in the respective input fractions (Figure 3A). To determine the efficiency and specificity precisely for individual transcripts, we spiked the 4sU-labeled total RNA from MCF-7 with *in vitro* transcribed 4sU-labeled FLuc and unlabeled RLuc that were followed by RT-qPCR analysis using a standard curve for quantification (Supplemental Figure 1). This analysis revealed a purification efficiency of 4sU-labeled FLuc of about 60% (58.56). The specificity was determined by the cross-contamination of RLuc in the biotin-enriched fractions and FLuc in the flow through fractions, which was about 5% for each transcript (RLuc in enriched = 5.32%, FLuc in flow through = 5.01%).

**Figure 3:**
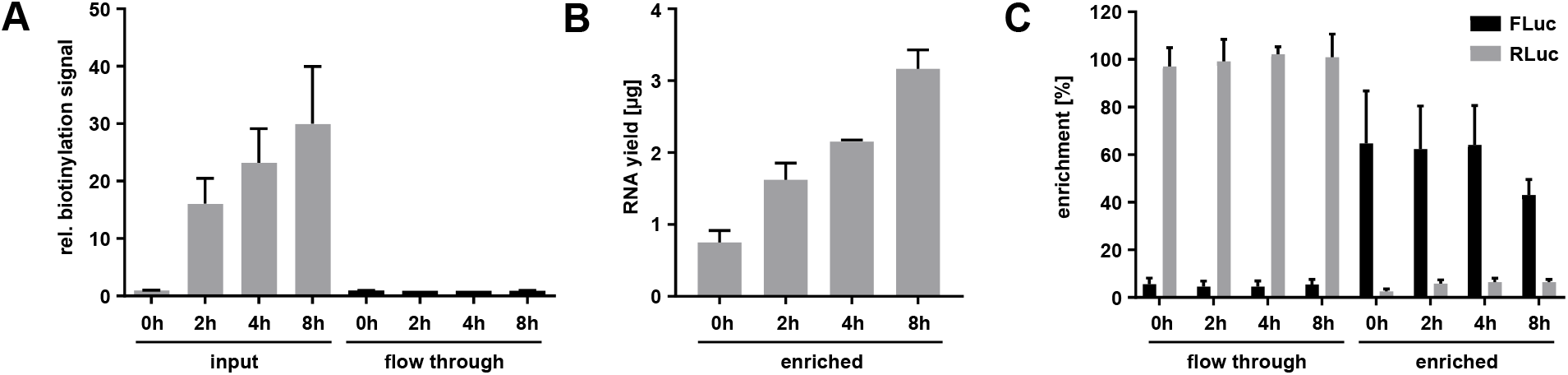
Purification of labeled and unlabeled RNA fractions. MCF-7 cells were pulse labeled with 4sU for up to eight hours as indicated. Total RNA was spiked with *in vitro* transcribed 4sU-labeled FLuc and unlabeled RLuc, biotinylated with MTSEA-biotin and subjected to streptavidin purification. (n = 3). **A:** Dot blot-based detection of biotinylation with streptavidin-HRP in input and flow through of streptavidin purification. **B:** The amount of RNA enriched by the streptavidin purification was determined by absorption measurement. **C:** *In vitro* transcribed spike in RNAs 4sU-labeled FLuc and unlabeled RLuc in the flow through and biotin-enriched fraction were measured by RT-qPCR analysis and normalized to a standard curve given in Supplemental Figure 1.

The kinetic model was fitted to the read counts from the sequenced samples for genes with mean read count *>* 50 in the total samples. Two total samples were collected at 0 hr, labeled and unlabeled fractions at other time points (2, 4 and 8 hrs) in two replicates (see Supplementary Table 1). In the model fitting, we assumed no cross-contamination between fractions and shared normalization coefficients for samples originating from the same time point and fraction.

Having the estimations for expression levels, degradation rates, overdispersion parameter and normalization coefficients, we calculated the FIM diagonal elements *ℐ*_dd_ for the analyzed genes for different time points and fraction types.

In Figure 4A, the value of the diagonal FIM element multiplied by *d̂*^2^, i.e. *ℐ*_dd_(***θ̂****, t*)*d̂*^2^ (compare to Equations 9 and 10), is depicted for both fractions. As mentioned in the previous section, *ℐ*_dd_ can be interpreted as an information gain from the experiment assuming other parameters were known, which represents an upper bound, see Equation 20. In addition, these terms are bounded due to presence of overdisperion in the data, (Equation 21 and dashed lines in Figure 4A), and increase of sequencing depth can not improve these limits.

**Figure 4:**
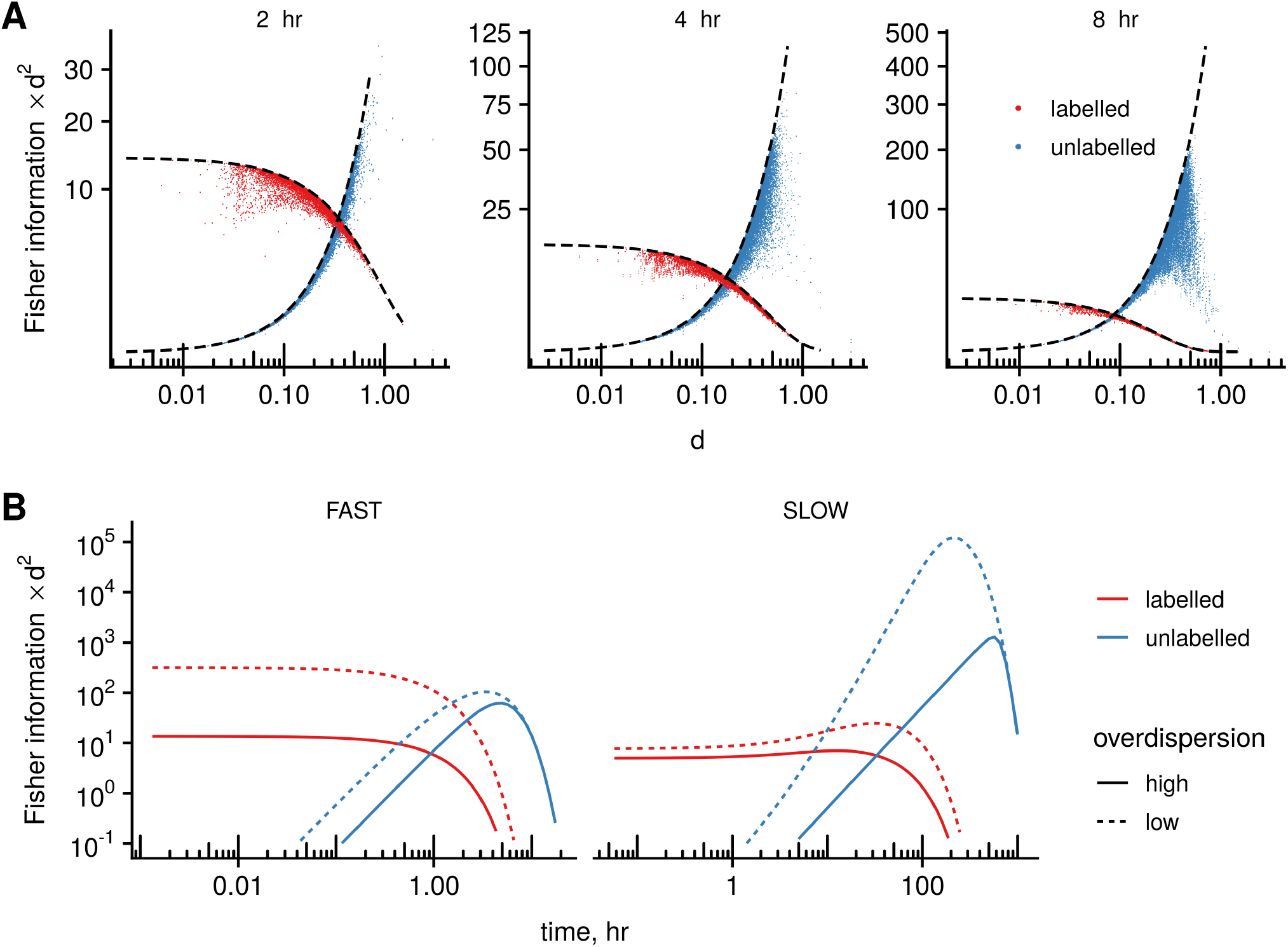
Application to experimental data from the MCF-7 pulse labeling time course experiment. **A:** We plot the diagonal term of the FIM computated at estimated parameter values and multiplied by *d̂*^2^, *ℐ* _dd_(***θ̂****, t*)*d̂*^2^, to illustrate contributions from labeled and unlabeled fractions to estimations of degradation rates for different experimental points (MCF7 experiment, 2, 4, and 8 hr) and fractions (labeled and unlabeled). The black lines are the limiting values for the *ℐ* _dd_ according to Equation 21. **B:** The modified FIM term *ℐ* _dd_(***θ̂****, t*)*d̂*^2^ is computed for a range of labeling times for one of the fastest (at the 0.1% quantile) and one of the slowest (at the 99,9% quantile) genes (*d*_fast_ = 0.79 hr^*−*1^, *d*_slow_ = 0.019 hr^*−*1^). The normalization coefficient is adjusted in such a way, that sequencing depth (total mean read count) at time *t* equals the sequencing depth of the total sample.

At short labeling times, the FIM term is higher for the labeled fraction than for the unlabeled one for majority of the genes, (Figure 4A, 2hr), which is a result similar to the SLAMseq case. At longer labeling times, the contribution from the unlabeled fraction increases, and (*ℐ*_U_)dd *>* (*ℐ*_L_)dd for majority of the genes (Figure 4A, 8hr). However, the proportion of genes with high degradation rates *d* in the unlabeled sample exponentially decreases, since

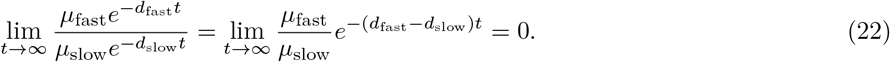

It results in very low counts and decrease in the *ℐ*_U_ for these fast genes, see Figure 4A, 8hr, reduced values at the right tail of the distribution (blue dots).

Optimal design for such experiments is complicated by the fact, that it depends not only on the degradation rates of some target genes, but on the overall rate distribution in the system being studied. We illustrate a dependency of the (*ℐ*)_dd_ *d*^2^ terms on labeling time for one of the fastest (0.1% quantile) and one of the slowest (99.9% quantile) genes. The normalization coefficients for labeled and unlabeled fractions were adjusted in the same manner as in the previous section, i.e. at every time point *t* the sequencing depth equals the sequencing depth of the total sample. In the case of low or no overdispersion, use of labeled fraction and shorter labeling times is preferred for estimation of fast genes, and, *vice versa*, longer labeling times and unlabeled fraction is preferred for slow genes, see Figure 4B, dashed lines.

At high values of overdispersion (i.e. low *k*), the FIM term is bounded (*ℐ*_L_)_dd_*d*^2^ *< k* due to Equation 21. In this case, there may exist values of labeling times at which the terms from the unlabeled fraction (*ℐ*_U_)_dd_*d*^2^ is larger than maximal (*ℐ*_L_)_dd_*d*^2^ value, Figure 4B, solid lines. As a protection against such situation in the case of fast genes, use of samples from unlabeled fraction may be a solution.

Although one may have a prior guess about the range of degradation rates in a system, it is unlikely, that there is information about the distribution of the rates and overdispersion level. Hence, such design suggestions are possible only in sequential approach, when an exploratory experiment is done first.

## Discussion

In this study, we discuss some aspects of the optimal design of RNA labeling experiments using the results of the asymptotic theory. First, we show that there exists an optimal time point for which the maximum likelihood estimator possess a minimal variance asymptotically. This first result was developed for the case of experiments, which preserve the fraction ratio and hence do not require normalization between fractions (e.g. SLAMseq, TUC-seq, TimeLapse-seq)

In the case of negligible overdispersion, the optimal labeling time for a gene with the characteristic degradation time *τ* is *t*_slam_ = 1.59*τ*, and shorter labeling times show better rate estimates in comparison to longer times: the variance increases exponentially for times longer than *τ* and only by a power law for shorter labeling times. This result is similar to the observations in a simulation study by Jürges et al. [2018]. Herein, for a given gene with half-life *λ* = 2 hr, the most precise estimation were at labeling times 3 hr and 6 hr (*t*_optimal_ = 1.59 2*/* log(2) = 4.6 hr), and the worst estimations were observed at the longest and the shortest times (12 hr and 0.5 hr). However, the exact ranking of time points is different for the given half-life time, probably due to influence of prior distribution utilized in the Bayesian framework.

We show that at short labeling times (in comparison to the characteristic time of degradation for a given gene), the labeled fraction contributes most to the Fisher information term corresponding to the degradation rate, and, *vice versa*, at long times the highest contribution is s een for the unlabeled one.

In addition, we show that in the presence of overdispersion, the variance of rate estimates is more sensitive to choices of labeling times different from the optimal, which make it more difficult to optimize conditions for a range of rates. The overdispersion imposes a bound on the asymptotic relative standard deviation for the estimator of the rate (*sd*(*d*)*/d*, see Figure 2C), and, from a certain level, increase in sequencing depth is very inefficient (Figure 2D).

Moreover, we discuss possible benefits of use of the conventional experimental approach, especially for estimation of extreme degradation rates, which deviate highly from the general pool. For nucleotide conversion setups with too short or too long labeling times, the majority of reads in a sample originate from the unlabeled or labeled fractions correspondingly. In contrast, the conventional scheme, which involves biochemical fraction separation, allows to concentrate experimental costs only on the relevant material.

Obviously, there are certain limitations to our study. First, the method involving FIM calculation describes only the asymptotic behavior of the estimator. Hence, all the conclusions are only approximate, since we do not investigate the behavior of the likelihood function itself, but only the quadratic approximation of its logarithm using the FIM.

Secondly, we do not consider uncertainty from the shared parameters, such as the overdispersion parameter of the negative binomial distribution and the normalization coefficients for the fractions. Inference on these parameters is based on the whole pool of the genes, and would involve more complex analytic treatment and assumptions on the distribution of rates.

Lastly, cross-contamination between fractions is a highly relevant problem for inference, especially in the absence of external reference molecules (spike ins), which are typically used to assess this phenomenon. However, in the Extended methods section, we show that cross-contamination shifts estimations of fast rates to slower values, and slow rates towards faster values. Previously, Eser et al. [2016] included a global transcriptome-wide cross-contamination term to presented kinetic model, yet future work is needed to assess possible effect sizes on rate estimations.

We hope that our work will encourage further development of the methodology to address the discussed limitations and to improve suggestions on design of metabolic labeling experiments.

## Acknowledgments

The authors thank Tobias Jakobi for computational infrastructure support and Etienne Boileau for providing useful comments on the paper draft. We are grateful for the excellent sequencing support by the Cologne Centre for Genomics. IND would like to acknowledge funding by the DFG (NA 1273/1-1). AU and CD acknowledge funding by the Klaus Tschira Stiftung gGmbH and DFG SPP 1784.

## Author Contributions

AU designed and performed the research. CD supervised the project. IND designed and performed the MCF7 experiments. AU, IND, CD analyzed the data. AU, IND and CD wrote the manuscript.

## Extended Methods

### Statistical model

We assume that the read counts follow the negative binomial distribution with the probability distribution function

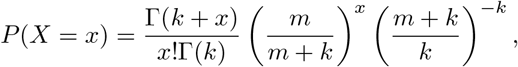

where *m* = *m*(*μ, d,…, t*) is the mean read count expected from the kinetic model and depends on the time point *t*, expression level in a steady-state *μ*, degradation rate *d*, and sample normalization, see the next section for details. The negative binomial distribution imposes a relation between variance and mean via the overdispersion parameter *k*, var*X* = *m*(*m* + *k*)*/k*. We assume the same *k* for all the genes in the data set, which is the simplest model, but more complicated models exist [Anders and Huber, 2012]. In this case, *k* is a shared parameter, and the model parameters from different genes must be fitted together in one procedure.

The logarithm of the likelihood function depends on the experimental points *X*_1_*,…, X_n_* and the vector of all model parameters ***θ***

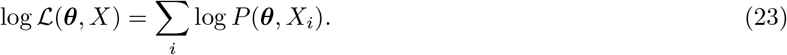

The maximum likelihood estimator is then

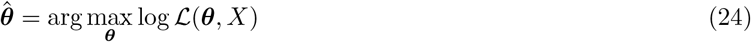

### Kinetic model

The solution of the differential equation for the kinetics of synthesis and degradation of the RNA 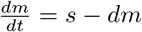 is

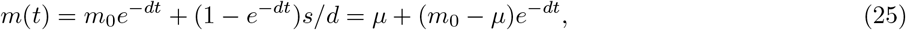

where *m*_0_ is the initial amount of RNA, *μ* = *s/d* is the expression level is the steady state for synthesis rate *s* and degradation rate *d*.

For definiteness, consider a pulse experiment. The unlabeled fraction is being only degraded (synthesis rate *s* = 0), and the initial RNA level is *μ*

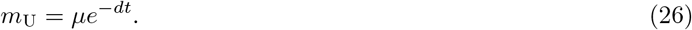

The labeled fraction starts from zero and saturates to the steady state:

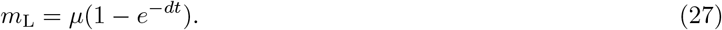

In fact, the mean read count is only proportional to the amount of RNA in samples, so without spike-in fragments added to measure absolute concentration, we can estimate the amounts only up to unknown coefficient. For identifiability, we use the read counts in the total samples as a reference (accounting for difference in sequencing depth, as implemented in the DESeq package [Anders and Huber, 2012]).

If the ratio of fractions is preserved, as, for example, in the SLAMseq protocol, the counts in the labeled *X*_L_ and the unlabeled *X*_U_ fractions are scaled by the same sequencing depth correction *x*, 𝔼(*X*_U_ + *X*_L_) = *xμ* and

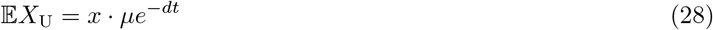

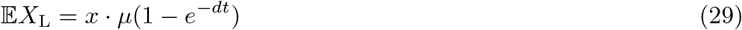

Assuming that usually samples are sequenced to approximately the same depth, for theoretical derivations we use *x* = 1 for the SLAMseq experiment.

If the fractions were separated by a chemical procedure, the read counts ratio will not coincide with the ratio of labeled and unlabeled molecules. In the case of negligible cross-contamination,

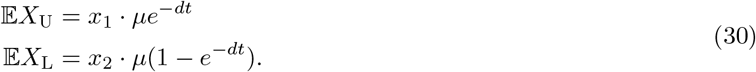

In this work, we do not consider cross-contamination, however it can play a significant role, especially for extreme rates (very fast or very slow) even if the overall contaminated material amount is low.

Indeed, if there is a cross-contamination level of *γ*, the mean read count can be modeled as

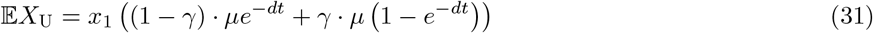

Under a wrong model, it leads to a biased degradation rate estimation:

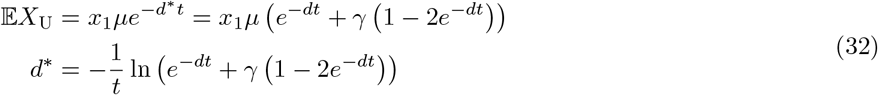

and for very slow genes, i.e. *dt* ≪ 1, using the first term of the Taylor expansion

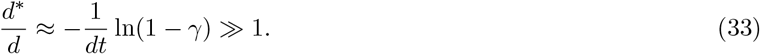

In contrast, for very fast genes, such as *dt* ≫ 1,

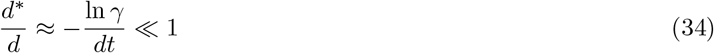

Hence, the rate estimations are biased to faster rates in the case of slow genes and toward faster values in the case of fast genes. The same result holds for the labeled fraction.

### Fisher information matrix derivation

Let *m*(***θ***, *t*) is the expected mean read number at time point *t* and ***θ*** = (*μ, d*)^*T*^is a vector of model parameters. The observed Fisher information matrix (FIM) is defined as

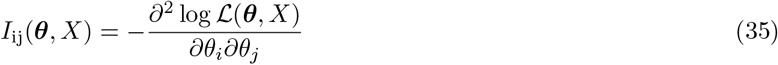

In the optimal design of experiments, the expected FIM is used

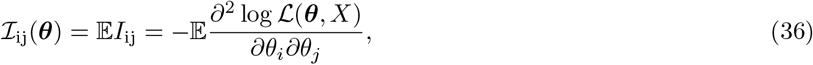

since we may not have any measurements and are interested in the performance of a given design in average. We refer everywhere only to the expected FIM and name it just FIM, omitting the “expected” term.

We will use here the fact, that the variance of the score function

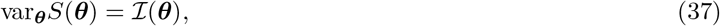

where

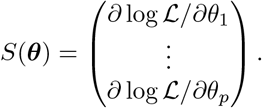

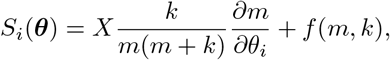

where *f* (*m, k*) is a term not depending on *X*. Using the fact that var*X* = *m*(*m* + *k*)*/k*,

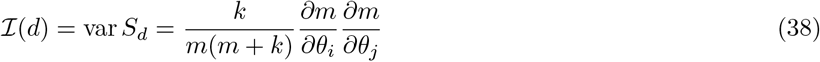

In the *k → ∞* case, when no overdispersion is assumed and the model follows the Poisson distribution,

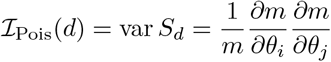

By plugging the relations for the RNA amount *m* = *μe^−dt^* (the unlabeled fraction) and *m* = *μ*(1 − *e^−dt^*) (the labeled fraction) in Equation 38,

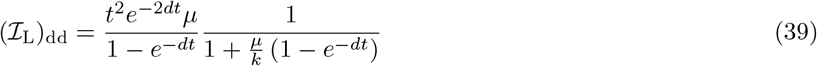

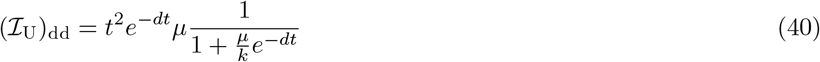

For the SLAMseq case,

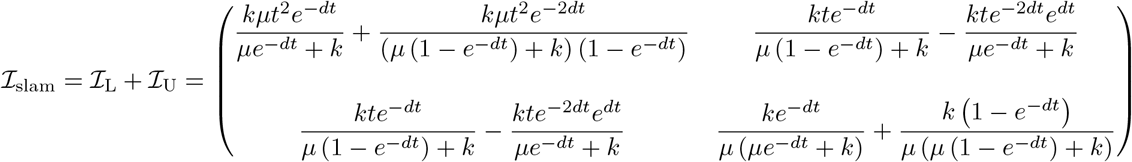

The inverse matrix in the general case is

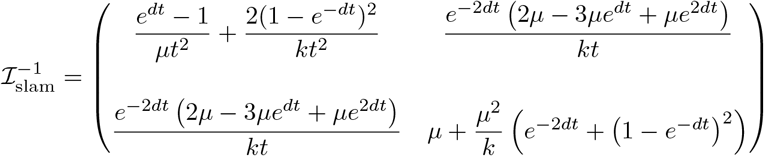

In the case of low overdispersion, *k* ≫ 1, it is simplified to

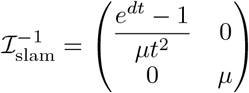

A model with overdispersion imposes a limit on the asymptotic lower bound of the MLE for *d*, which cannot be improved by increase of sequencing depth (i.e. *μ*). For a fixed *t*,

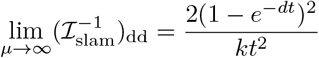

Although this limit value decreases with *t* →∞, the depth *μ* must increase exponentially *e^dt^/μ*(*t*) → 0 in order to satisfy

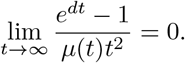

### Normalization in conventional experiments

The situation is different for the case of biochemical purification, when the labeled, unlabeled and total fractions are sequenced to the approximately same depth. For the next derivation, we assume that samples were normalized externally, e.g. by synthetic spike-ins or exogenous RNA. In addition, we do not consider uncertainty coming from fraction normalization.

The sequencing depth for the total sample is Σ_*i*_μ_*i*_. After labeling for *t* hours, the concentrations of labeled and unlabeled molecules changes according to Equation 30, and in the case of same depth, the normalization coefficients are

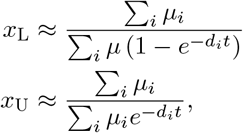

where we use the total sample as a reference, i.e. with the normalization coeffiecient 1. At long times, when majority of genes in the labeled fraction achieve saturation, *x*_L_ ≈ 1. Similarly, at short times, *x*_U_ ≈ 1, degradation has a minor effect on the unlabeled fraction.

In contrast, at short times,

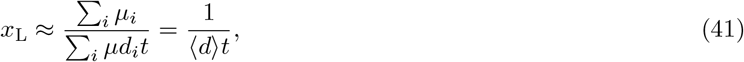

where 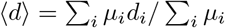 is average degradation rate weighted by the steady-state expression level. If there is a small cluster of fast genes *i* ∈ *ℱ*, which dominate the pool of labeled molecules at such times, that 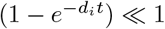 for other genes (*i* ∉ *ℱ*), and 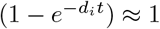 for *i* ∈ *ℱ*, then

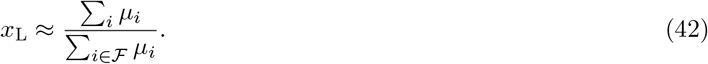

Similar result holds true for the cluster of slow genes *S*, such that 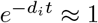 for *i* ∈ *S* and 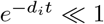 for *i* ∉ *S*, and

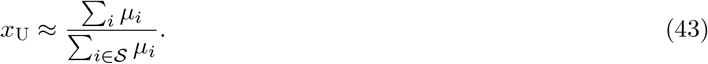

Such “zooming” effect of normalization can drastically improve inference about fast genes in comparison to the SLAMseq design, since *x*_L_ and *x*_U_ can be interpreted as a corresponding increase in depth in SLAMseq experiment.

Modifying the mean read count in Equations 39 and 40 by the factors *x*_L_ and *x*_U_ results in

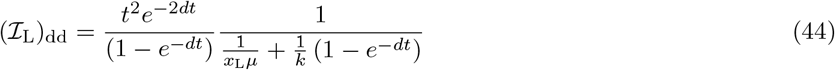

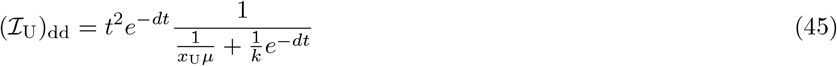

As in the SLAMseq case, the overdispersion imposes limits on the terms of the FIM:

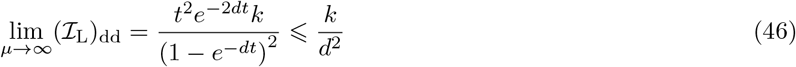

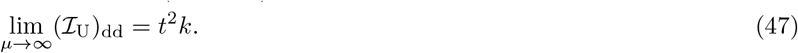

It is interesting, that in the case of the unlabeled fraction, this upper bound can be improved by using longer labeling times, which is not true for the labeled one.

## Experimental Model and Methods

### Tissue Culture Cell Line

MCF-7 cells (ACC-115) were obtained from the Leibniz Institute DSMZ German Collection of Microorganisms and Cell Cultures. Cells were routinely tested for mycoplasma contamination with Venor GeM Classic (Minerva Biolabs). MCF-7 cells were cultured at 37°C and 5% CO2 and maintained in DMEM (Thermo Fisher Scientific) supplemented with 10% fetal calf serum (Merck), 1xMEM non-essential amino acids (Thermo Fisher Scientific) and 1xPenicillin/Streptomycin (Thermo Fisher Scientific).

### Tissue Culture

MCF-7 cells were seeded 48 hrs prior to the experiment at a cell density of 0.3 105cells*/*cm2. Cells were labeled with 4-thiouridine (4sU) (Sigma-Aldrich) at a final concentration of 200 *μ*M for 2, 4 or 8 hrs. Cells were scraped in DPBS and the pellet resuspended in Trizol (Thermo Fisher Scientific).

### Isolation of total RNA

Total RNA was isolated using the Trizol method. Briefly, the cell pellet was resuspended in 750 *μ*l Trizol, and incubated 5 min at room temperature before addition of 200 *μ*l chloroform. Samples were centrifuged (20 min, 10.000g, room temperature) and the aqueous phase re-extracted with one volume chloroform: isoamylalkohol (24:1) (5 min, 10.000g, room temperature). The RNA in the aqueous phase was precipitated with one volume isopropanol (30 min, 20.8000g, 4°C), washed twice with 1 ml 80% ethanol in DEPC-H_2_O and dissolved in 25 *μ*l DEPC-H_2_O (10 min, 55°C, shaking).

### *In vitro* transcription of spike ins

For *in vitro* transcription of linearized plasmids (pBSIIKS-Luc-pA-NB [Liepelt et al., 2014] and pBSIIKS-Renilla-pA [Thermann and Hentze, 2007]), the MEGAscript T7 Transcription Kit (Thermo Fisher Scientific) was used according to the manufacturers instructions. Briefly, the reaction was set up in a total volume of 20 *μ*l containing 1 *μ*g linearized plasmid and 2 *μ*l 10x reaction buffer, 3 *μ*l 40 mM m7GppG-cap analogon (KEDAR), 2 *μ*l 15 mM GTP, 2 *μ*l 75 mM CTP, 2 *μ*l 75 mM ATP, 2 *μ*l enzyme mix and 2 *μ*l 75 mM UTP (for RLuc) or 2 *μ*l 75 mM 4-S-UTP:UTP in a 1:10 ratio (for FLuc). Reactions were incubated 3 hrs at 37°C. Plasmid-DNA was removed by addition of 1 *μ*l Turbo-DNase (15 min, 37°C). *In vitro* transcribed RNA was purified by phenol extraction and Chromaspin-100 (Clontech) purification. RNA was precipitated over night after addition of sodium acetate to a final concentration of 0.3 M and 2.5 volumes 100% ethanol. After centrifugation (30 min, 20.800g, 4°C) the pellet was washed with 1 ml 80% ethanol and dissolved in 40 *μ*l DEPC-H_2_O. Concentration was determined by Nanodrop (Thermo Fisher Scientific) measurement and integrity checked by agarose gel electrophoresis.

### Biotinylation of RNA

Total RNA was spiked with *in vitro* transcribed 4sU-labeled FLuc and non-labeled RLuc RNAs and biotinylated using MTSEA biotin-XX (Biotium) as described by [Duffy et al., 2015]. Briefly 80 *μ*g total RNA was incubated with 8 ng FLuc and 4.8 ng RLuc (equimolar amounts, 130 amol), 10 mM HEPES pH 7.5, 1 mM EDTA and 5 *μ*g MTSEA biotin-XX (freshly dissolved in DMF) in a total volume of 250 *μ*l. Reactions were incubated 30 min in the dark at room temperature. Biotinylated RNA was recovered by extraction with one volume phenol: chloroform: isoamylalkohol (24:24:1) and separated using Phase-Lock-tubes (5Prime) by centrifugation (5 min, 20.800g, room temperature). RNA was precipitated by addition of 350 *μ*l isopropanol, 25 *μ*l 5 M sodium chloride and 1 *μ*l glycogen (Roche Diagnostics, 20 *μ*g/*μ*l) to assist precipitation (30 min, 20.800g, 4°C). RNA was washed twice with 500 *μ*l 80% ethanol in DEPC-H_2_O and dissolved in 25 *μ*l DEPC-H_2_O (10 min, 55°C, shaking).

### Streptavidin purification

For purification of biotinylated RNAs the method described by [Schwanhäusser et al., 2011] was adapted. 25 *μ*g biotinylated total RNA was adjusted to 100 *μ*l with DEPC-H_2_O and filled up with Streptavidin binding buffer (Strep-BB) (20 mM Tris, pH 7.4, 0.5 M sodium chloride, 1 mM EDTA) to 200 *μ*l. RNA was denatured 10 min at 65°C and subsequently placed on ice. 100 *μ*l magnetic streptavidin beads (New England Biolabs) were washed once with 200 *μ*l Strep-BB and resuspended in 100 *μ*l Strep-BB. RNA and beads were incubated 15 min at room temperature on a rotating wheel. Beads were washed three times with 500 *μ*l Strep washing buffer (100 mM Tris pH 7.4, 1 M sodium chloride, 10 mM EDTA, 0.1% Tween 20) prewarmed to 55°C. RNA was eluted three times by de-biotinylation with 100 *μ*l freshly prepared 100 mM DTT and elution fractions pooled for further analysis. RNA was recovered from total RNA, flow through and eluate by phenol: chloroform: isoamylalkohol (24:24:1) extraction using Phase-Lock-tubes and isopropanol precipitation as described above. The amount of recovered RNA was determined by Nanodrop measurement.

### Dot blot-based detection of biotinylation

1 *μ*g biotinylated RNA was applied to nylon membrane (Hybond-N, GE Healthcare) using a dot blot device (Carl Roth). RNA was crosslinked twice at 254 nm using the "Optimal Crosslink" mode of the Spectroline Select XLE-1000 crosslinker. The membrane was blocked 20 min with PBS + 10% SDS and incubated 2 hrs with Streptavidin-HRP (Thermo Fisher Scientific, 1:5000 in PBS + 10% SDS). Prior to detection with SuperSignal West Pico (Thermo Fisher Scientific) the membrane was washed each three times 10 min with PBS + 10% SDS, PBS + 1% SDS and PBS + 0.1% SDS. Images were acquired with the LAS4000 system (GE Healthcare).

### Reverse Transcription

1 *μ*l RNA from streptavidin purification was reverse transcribed using the Maxima H Minus First Strand cDNA Synthesis Kit (Thermo Fisher Scientific) with Random Primers according to the manufacturers protocol. For absolute quantification reverse transcription reactions were set up with different amounts of spike in RNAs, ranging from 1600% to 1.56% for FLuc and 400 to 3.12% for RLuc in 1:2 dilutions. Briefly, RNA was mixed in a total volume of 15 *μ*l with 1 *μ*l Random Primer and 1 *μ*l dNTP solution and denatured (5 min, 65°C). Reaction was completed by addition of 4 *μ*l 5xRT buffer and 1 *μ*l Maxima enzyme and incubated 10 min at room temperature followed by 30 min, 50°C and denaturation (5 min, 85°C).

### qPCR Analysis

Reverse transcription reactions were diluted 1:10 and used for qPCR analysis on a StepOnePlus instrument (ThermoFisherScientific) with Power SYBR Green PCR Master Mix (Thermo Fisher Scientific) and primers directed against FLuc (forward: CCTTCCGCATAGAACTGCCT, reverse: GGTTGGTACTAGCAACGCAC [de Vries et al., 2013]) and RLuc (forward: GTTGTGCCACATATTGAGCC, reverse: CCAAACAAGCACCCCAATCATG [Naarmann-de Vries et al., 2016]).

### Sequencing

Total and enriched samples were depleted for ribosomal RNA (rRNA) contamination using RiboZeroGold, which is based on the removal of rRNA with biotinylated oligos using streptavidin beads. Thus, also the biotinylated 4sU-labeled molecules were removed from the total samples by the RiboZeroGold procedure and were treated as flow through. Libraries of 2 biological replicate 4sU pulse experiment were sequenced 1x 50bp on an Illumina HiSeq4000. All relevant details on sequencing depth and mapping rates are listed in Supplementary Table 1.

### Read processing and counting

Sequencing adapters and low-quality reads were removed from the raw sequencing data with flexbar v3.0.3 [Roehr et al., 2017] using standard filtering parameters. We excluded all reads with more than 1 uncalled base from the output. All remaining reads (>18bp) were then aligned to a custom sequence index including rRNA, tRNA and snoRNA gene loci using bowtie2 with the –very-fast option [Langmead and Salzberg, 2012]. Only reads that did not align to any of the contaminant sequences were considered for further analysis.

Reads were then aligned to the human genome (EnsEMBL 85) and splice sites from the reference annotation with a splice-aware aligner (STAR, v2.5.3a; Dobin et al. [2013]). The BAM files were analyzed with StringTie 1.3.3b [Pertea et al., 2015] and the final read count matrix was prepared with the supplemented python script prepDE.py.

## Supplementary Information

**Figure S1:**
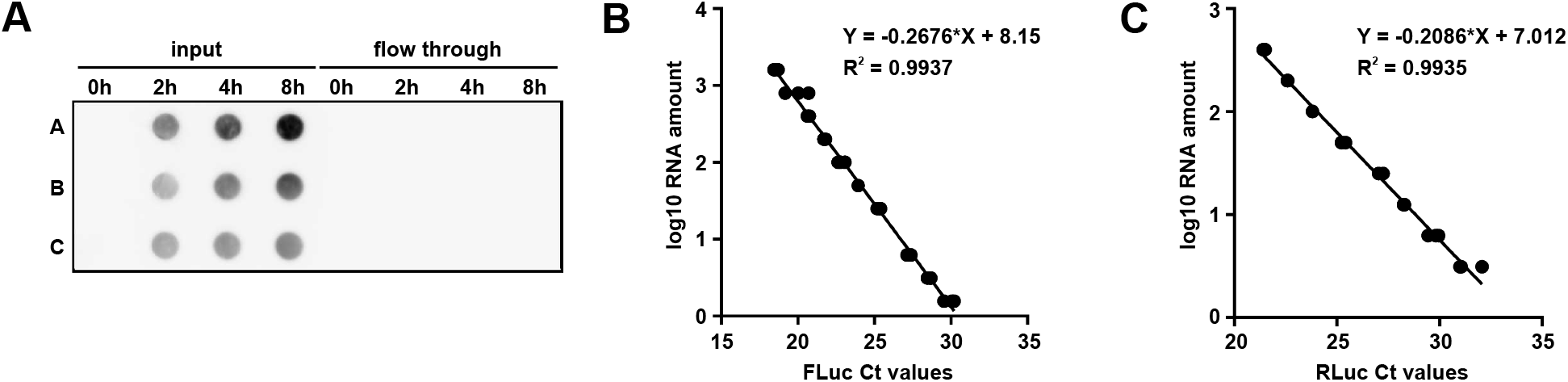
**A:** Dot blot-based detection of biotinylation with streptavidin-HRP in input and flow through of streptavidin purification from three replicate experiments A-C. The quantification of the captured image is shown in Figure 3A. **B:** Standard curve for the absolute quantification of 4sU-labeled FLuc RNA. 1600 to 1.56% of the input used for streptavidin purification was measured by RT-qPCR analysis in 1:2 dilutions. The log10 amount of RNA was plotted against the obtained Ct value and used for linear regression. **C:** Standard curve for the absolute quantification of unlabeled RLuc RNA. 400 to 3.13% of the input used for streptavidin purification was measured by RT-qPCR analysis in 1:2 dilutions. The log10 amount of RNA was plotted against the obtained Ct value and used for linear regression.

**Table S1:** Summary of RNA-seq read mapping statistics.

| Condition | Sample ID | Uniquely mapped reads % | Uniquely mapped reads number |
| --- | --- | --- | --- |
| Rep2 8h pull-down | K002000165_76894 | 90.4% | 31.4 |
| Rep2 0h Total RNA | K002000165_76874 | 85.1% | 29.5 |
| Rep1 8h pull-down | K002000165_76872 | 89.1% | 28.2 |
| Rep2 2h pull-down | K002000165_76882 | 92.5% | 27.3 |
| Rep1 4h pull-down | K002000165_76866 | 89.3% | 26.8 |
| Rep2 4h pull-down | K002000165_76888 | 91.2% | 24.2 |
| Rep1 0h Total RNA | K002000165_76852 | 79.9% | 23.2 |
| Rep1 2h pull-down | K002000165_76860 | 91.3% | 23.2 |
| Rep1 2h flow-through | K002000165_76856 | 73.7% | 21.8 |
| Rep2 2h flow-through | K002000165_76878 | 77.1% | 21.7 |
| Rep2 4h flow-through | K002000165_76884 | 71.5% | 21.0 |
| Rep2 8h flow-through | K002000165_76890 | 66.0% | 18.0 |
| Rep1 8h flow-through | K002000165_76868 | 63.4% | 16.6 |
| Rep1 4h flow-through | K002000165_76862 | 66.2% | 11.4 |

Table S2: **Read counts for all samples Weblink**

